# High Throughput Tomography (HiTT) on EMBL Beamline P14 on PETRA III

**DOI:** 10.1101/2023.09.06.556509

**Authors:** Jonas Albers, Marina Nikolova, Angelika Svetlove, Nedal Darif, Matthew J. Lawson, Thomas R. Schneider, Yannick Schwab, Gleb Bourenkov, Elizabeth Duke

## Abstract

Here we present High-Throughput Tomography (HiTT), a fast and versatile phase-contrast imaging platform for life-science samples on the EMBL beamline P14 at DESY in Hamburg, Germany. We use a high photon flux undulator beamline to perform tomographic phase contrast acquisition in about two minutes which is linked to an automated data processing pipeline that delivers a 3D reconstructed data set less than a minute and a half after the completion of the X-ray scan. Combining this workflow with a sophisticated robotic sample changer enables the streamlined collection and reconstruction of X-ray imaging data from potentially hundreds of samples during a beamtime shift. HiTT permits optimal data collection for many different samples and makes possible the imaging of large sample cohorts thus allowing population studies to be attempted. We demonstrate the successful application of HiTT on various soft tissue samples in both liquid (hydrated and also dehydrated) and paraffin embedded preparations. Furthermore, we demonstrate the feasibility of HiTT to be used as a targeting tool for volume electron microscopy (vEM), as well as using HiTT to study plant morphology. We also show how the high throughput nature of the work has allowed large numbers of “identical” samples to be imaged to enable statistically relevant sample volumes to be studied.

**Synopsis:** We present HiTT – high throughput tomography – a propagation based phase contrast X-ray imaging technique which can visualise 1 mm^3^ biological samples of various types at high resolution. The 3D reconstructions of the imaged volumes are calculated automatically once data collection is complete. The entire process from pressing start on data collection to viewing the final data takes less than 3 minutes. This speed in combination with the use of the automated sample changer to exchange the samples truly enables high throughput X-ray imaging for the first time.

## 1. Introduction

High resolution X-ray nanotomography is becoming an increasingly popular tool for the 3D analysis of samples in the life sciences (Albers, Pacilé *et al*., 2018; Eckermann *et al*., 2020; Frohn *et al*., 2020; Dahlin *et al*., 2020; Palermo *et al*., 2022). Using state of the art laboratory systems, both commercial and scientific, the acquisition of submicron resolution data sets has become possible, but micro-/nano-focus X-ray tubes come with the caveat of a relatively low X-ray photon flux, which in turn leads to scan times of multiple hours (Rawson *et al*., 2020; Albers, Markus *et al*., 2018). In addition, biological samples such as tissues, organs or cells have a very low X-ray absorption which leads to a lack of contrast for X-ray imaging applications. To circumvent this issue in laboratory-based X-ray imaging the go to method is the use of different staining protocols based on heavy metal ions such as tungsten or osmium to increase soft tissue contrast (Dullin *et al*., 2017; Metscher, 2009*a*,*b*). These staining techniques can be rather time consuming and/or complicated and create an artificial contrast. Additionally, uniform take up (absorbance) of the stain in large samples is often a challenge. An alternative approach, enabling the increased impact of X-ray imaging in life-science research, is the application of phase contrast which can generate far greater image contrast compared to absorption-based imaging (Endrizzi, 2018; Kitchen *et al*., 2017; Saccomano *et al*., 2018). This is especially true for low X-ray absorbing samples such as soft tissues (Momose *et al*., 1996; Nugent, 2010). In propagation-based phase-contrast imaging contrast is generated by the transformation of the phase shifts generated as the X-rays pass through the sample into measurable intensity variations by self-interference in free space propagation between sample and detector (Paganin & Nugent, 1998; Cloetens, Ludwig, Baruchel, Guigay *et al*., 1999). This requires a certain degree of spatial coherence which is offered by synchrotron light sources and, to a lesser extent, nano-focus X-ray tubes (Otendal *et al*., 2008; Töpperwien *et al*., 2017). The greater the degree of coherence of the incoming X-ray beam, the greater the amount of contrast that can be generated. Other phase contrast methods are possible such as edge illumination and/or grating based techniques. However to date the easiest way to collect data using phase contrast is by using synchrotron X-rays.

Even though synchrotron-based biological X-ray imaging has come a long way since its conception roughly 20 years ago, imaging experiments still lack the streamlined pipelines that are present for example in macromolecular crystallography (MX) (Oscarsson *et al*., 2019). While the use of synchrotron facilities can reduce the need for hour-long acquisition times, faster scanning results in a very high raw-data output which non-expert users in particular might not be able to handle. Consequently the scientific output of precious synchrotron scan time is often less than expected.

Now we want to take the experience gained from MX with its well established, state-of-the-art beamline infrastructure and apply it to phase contrast X-ray imaging. The first steps in this process involved the use of phase contrast imaging to visualise protein crystals (Polikarpov *et al*., 2019).

## 2. Methods

### 2.1. Beamline Overview

All experiments were carried out on the EMBL beamline P14 at the PETRA III storage ring, DESY, Hamburg, Germany. P14 uses a standard U29 undulator (Tischer *et al*., 2007; Key Parameters of Undulators Operated at PETRA III) with a nominal source size of approximately 13x330 μm (FWHM) in the vertical and horizontal directions, respectively. Employing a liquid-nitrogen-cooled vertical offset double-crystal Si(111) monochromator, P14 can operate with X-ray energies ranging from 6 to 30 keV. For X-ray imaging, energies between 10 keV and 27 keV have been successfully used. The beamline layout is shown in Fig. 1a. While for crystallographic experiments the beam size and shape of the X-ray beam can be adjusted by using refractive and/or reflective optical elements, for X-ray tomography experiments an ‘unfocused’ configuration is used in which neither compound refractive lenses nor X-ray mirrors interfere with the beam. Switching between the imaging and conventional MX optical arrangements can be done in a few seconds via the click of a button in the graphical user interface, MXCuBE which controls the beamline (MXCuBE v2 QT; (Gabadinho *et al*., 2010; Oscarsson *et al*., 2019); experimental parameters and intermediate results are stored in the attached ISPyB database (Delagenière *et al*., 2011).

**Figure 1.**
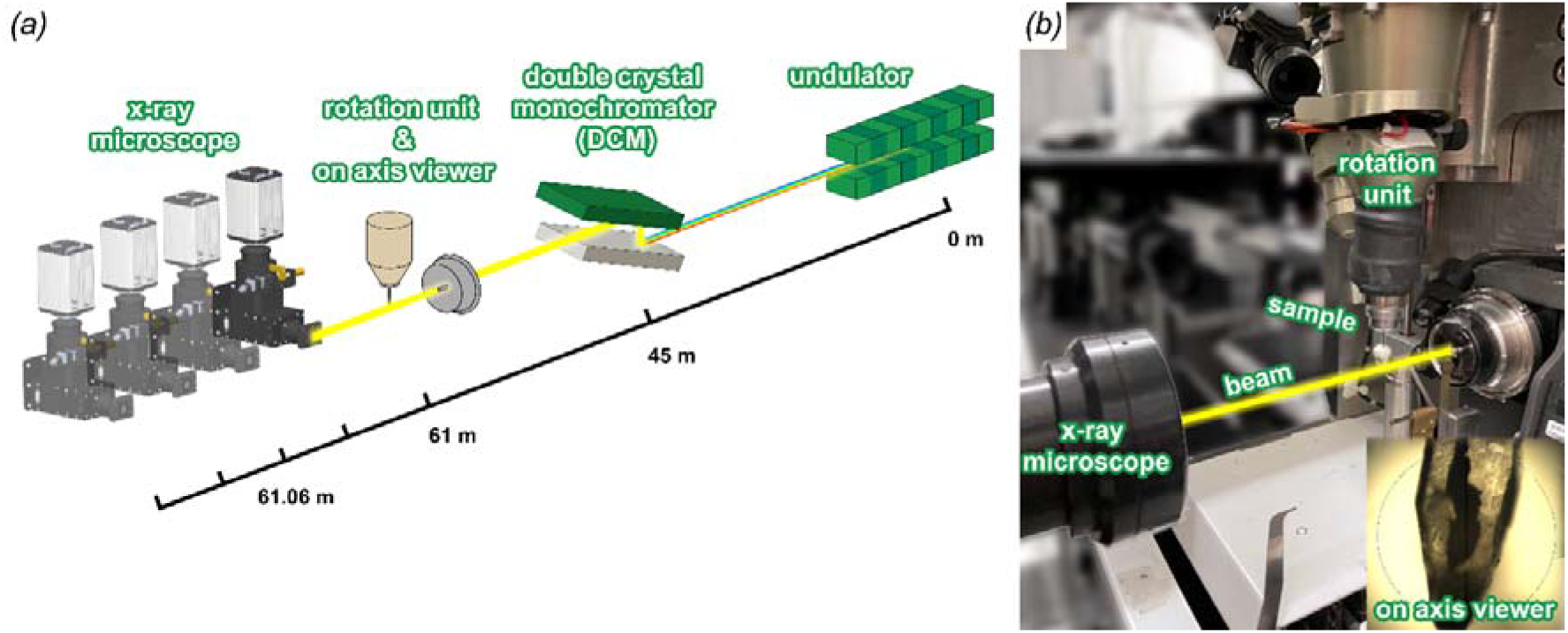
HiTT setup for nano-holotomography EMBL Beamline P14 on PETRA III.

For sample translation and rotation a modified MD3 diffractometer (Arinax, Moirans, France; (Cipriani *et al*., 2007)) is used. In contrast to common X-ray imaging setups the samples are suspended from above. Sample rotation is realised with a <100 nm sphere of confusion for the vertical and downward axis combined with the sample-centring stage. The MD3 houses an on-axis viewing system (OAV) with a zoomable optical microscope which can be operated in both transmission and reflective mode. The OAV is used for sample positioning and alignment.

The in-house designed detector stage supports an EIGER2 X CdTe 16M (DECTRIS, Baden, Switzerland) for crystallographic measurements as well as an X-ray imaging microscope (Optique Peter, Lentilly, France). The stage offers five degrees of freedom (vertical and horizontal translation, roll, pitch and yaw). The sample-to-X-ray imaging microscope distance is adjustable between 6 cm and 3 m. The high photon flux of the 6 GeV PETRA III storage ring on P14 (10^13^photons/s per 0.666x0.666 mm^2^ at 13 keV with a tuned undulator) saturates the detector within the X-ray microscope so for the tomography experiments the undulator is detuned by altering the undulator gap in order to reduce the flux by a factor of between 2 and 4.

### 2.2. Imaging Setup

X-ray images are obtained using a monochromatic X-ray beam imaging microscope (Optique Peter, Lentilly, France) equipped with an LSO:Tb scintillator with a thin active layer of 8 μm (ESRF, Grenoble, France), an Olympus UPlanFL 20-fold objective lens (Olympus, Tokio, Japan), a 45°-reflecting mirror, an Olympus 180 mm tube lens, and a PCO.edge 4.2 sCMOS camera (PCO, Kelheim, Germany) with a 2048×2048 pixel sensor with a pixel size of 6.5 μm. The effective pixel size of 0.325 μm results in a 666×666 μm^2^ field of view (FOV). For tomographic imaging in the near-field edge-enhancing regime four single tomograms at increasing sample to detector distances are acquired. For each distance, 1810 projections over a 181° rotation angle are recorded using an exposure time of t_exp_=10 ms and a frame-rate of 100 Hz resulting in a total exposure time of 4×18.1 s = 72.4 s. In addition 100 flat-field frames are recorded per distance. Sample positioning and alignment for tomography is performed using a “3-click-centring” approach developed for MX utilising the on-axis viewing optical microscope thereby removing the requirement to use X-rays for sample positioning (Gabadinho *et al*., 2010). As a result the incident X-ray dose on the sample is solely reduced to the exposure time required for data acquisition. A photograph of the X-ray imaging setup is shown in Fig. 1b.

For performing tiled acquisitions the sample is initially positioned via “3-click-centring” and then further moved by a predefined distance while maintaining a constant start-angle for the sample rotation for each data acquisition.

### 2.3. Sample Mounting and Handling

Unified sample mounts between different synchrotrons and/or beamlines are the norm in macromolecular crystallography. We are taking advantage of these developments by repurposing the existing macromolecular crystallography infrastructure at P14 for X-ray imaging.

HiTT samples are mounted on the MD3 diffractometer using magnetic “SPINE-style” goniometer bases (such as B5 goniometer base, MiTeGen, Ithaca, USA; (Cipriani *et al*., 2006)).

In order to enable sample mounting using the automated sample mounting system (MultiAxesRoboticVersatileINstaller, MARVIN; (Ristau *et al*., 2019)) the total dimensions of the sample plus mount must conform to the established dimensions in order to ensure reliable mounting and dismounting (max sample length including pin of 25 mm, max sample diameter 8 mm). To date samples measured tend to fall into the following categories (Fig. 2a): samples embedded in resin, samples embedded in paraffin wax and samples mounted in liquid. Resin-embedded samples can be glued to the tip of a pin in the same way that a crystal mounting loop or mesh is attached for structural biology data collection. Alternatively, custom-made 3D printed tips have been designed which can be attached to the spine base. The polylactide plastic material in combination with the slightly wider tip end enables easier gluing of samples which is particularly helpful for larger width samples such as 1 mm cylinders of paraffin-embedded tissues. Samples imaged in liquid are mounted in a modified, cut down 10 μl pipette tip. To date we have imaged samples suspended in both PFA (1-4% in buffer) and ethanol. In our hands we find, particularly for tissue samples, that dehydration in ethanol results in slightly higher quality data due to increased contrast. Sample stability during imaging is helped by the fact that the sample is suspended from the MD3 rather than mounted from below. The 1 mm tissue punch biopsy core sits nicely in the tip of the pipette enabling imaging without sample movement. The challenge is to ensure that no bubbling occurs during data collection and that, if it does take place, it does not result in sample movement so that the data quality is not compromised. We find that fewer bubbles are generated if the liquid surrounding the samples is ethanol rather than an aqueous solution. We found that for a kidney sample mounted in this manner we were able to collect high quality data on a sample that had been sitting on the bench in its mount for 6 months suggesting that this method of sample mounting does not compromise tissue quality.

**Figure 2.**
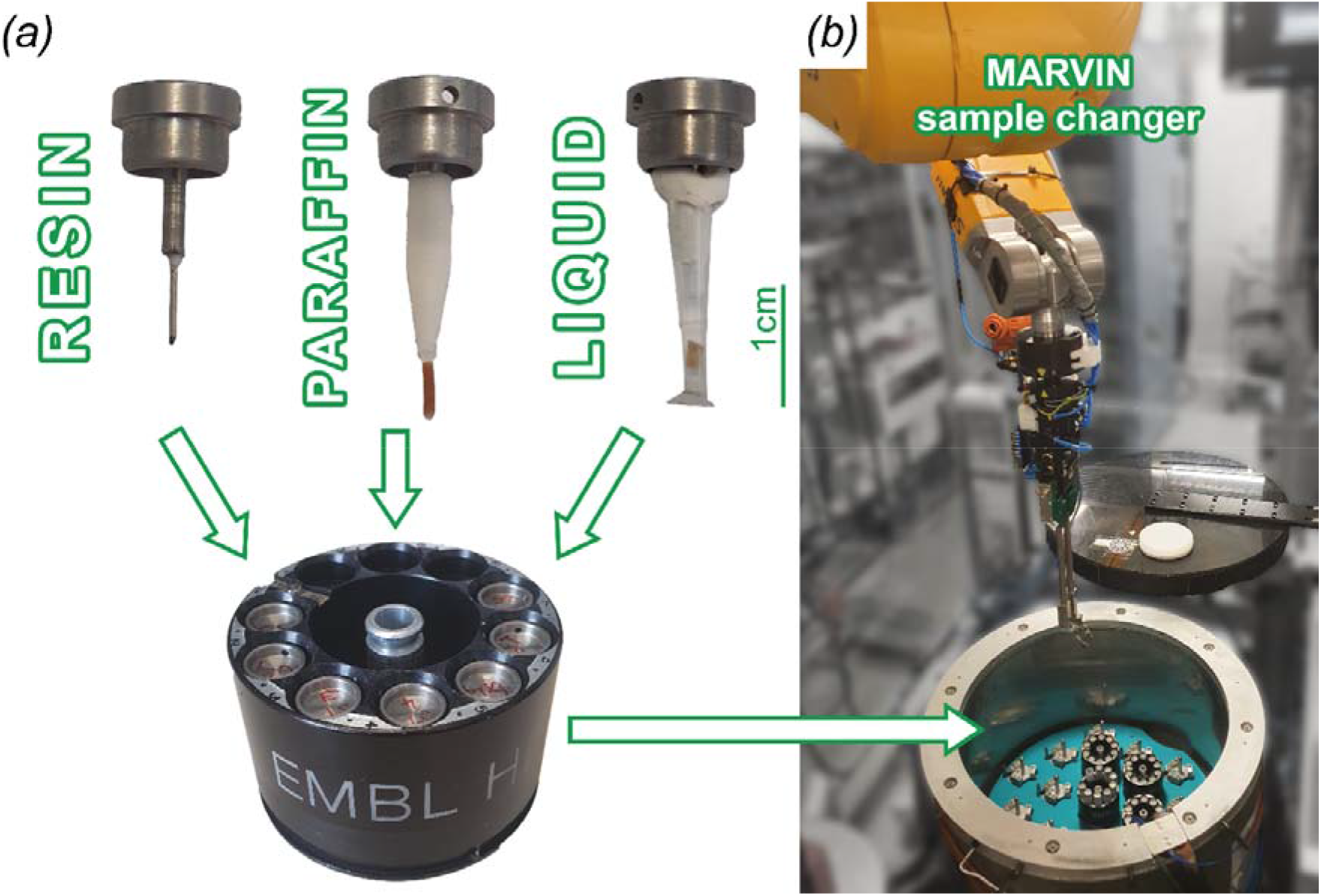
Sample handling at the P14 beamline Samples are mounted on SPINE style pins. Mounting schemes for resin embedded, heavy-metal stained samples, paraffin embedded, as well as wet samples mounted in liquid were established. SPINE pucks hold up to ten samples each. Up to 17 spine pucks (=170 samples) can be loaded into the MARVIN sample changer. Samples are mounted using a robotic arm with a dedicated sample gripper.

Mounted samples are loaded into SPINE pucks, metal carriers which can hold 10 samples at a time. These are then loaded into the MARVIN dewar, which has a capacity of 17 pucks (=170 samples). When the beamline is used in “MX-mode” the dewar is filled with liquid nitrogen for measurements at cryogenic conditions. In contrast, HiTT imaging is carried out at room temperature. This does not affect the performance of MARVIN in any way.

### 2.4. Data Acquisition and Processing

Data are normally acquired on an external trigger synchronised with the rotation angle of the diffractometer. An in-house developed PCO CAMERA server (C/C++) communicates directly with the camera, preparing it for data acquisition given the user-provided parameters and setting it up in a wait for the hardware trigger signal. The frames are extracted continuously, stored in pre-allocated buffers, and sent out over a 40 Gbit/s InfiniBand connection and a Zero MQ (https://zeromq.org/) stream to the ARAPAHO server in raw format.

ARAPAHO, another in-house development (C/C++), has several main tasks – (a) receive the raw data over the stream, (b) for each of the projections find the best matching background/flat-field frames using a correlation algorithm optimised for speed, (c) flat-field correct a subset of the projections by their respective best-matching background frame, (d) compress this subset into 8-bit jpegs, (e) send out the jpegs to a live viewer of flat-field-corrected frames, and (f) dump the raw data converted to tiff format to disk. Those tasks are executed in parallel with the data acquisition.

The correlation (task (b) above) is applied over a predefined region of interest, by default set in the PCO CAMERA server to 600x1850 pixels with x, y coordinates 300x100 to 900x1950. This default can be modified so that a different area of the 2048x2048 image is chosen – in order to exclude unwanted artefacts on the image, for example. Computing the correlation of each projection with 100 background frames over an area of more than 600 pixels in the horizontal direction at maximum or close to maximum pixel height is, however, not possible at the standard acquisition rate of 100 Hz as then ARAPAHO would no longer be able to keep up. Such a large region of interest, while nice to have for some samples and experimental conditions, does not seem to be a requirement for good final reconstructions.

Over a data collection of 1810 frames, 423 frames are typically flat-field corrected with their best-matching background frame (task (c)). Those 423 frames are spread equidistantly across the range of 1810 frames according to the following condition, flat-field correct a given frame if

*n t*_*exp*_*-floor(n t*_*exp*_*22) / 22* ≤ *t*_*exp*_ where n is the frame number. This results in about 23 flat-field-corrected frames per second which are then normalised in order to fit the 16-bit raw data pixel sizes into the 8 bits used by the jpeg compression (task (d)). The jpegs are sent out over a 1 Gbit/s connection, via the TINE control system (Bartkiewicz & Duval, 2007; Duval *et al*., 2018), and the TCP/IP network protocol (task e).

When creating the tiffs in which we store the data to disk to each projection file we add a couple of custom tiff tags. The first contains the indexes of the 3 best-matching background frames, and the second a list of 100 correlation coefficients, the result of the computation of the correlation between the projection frame and each of the 100 background frames.

The tiffs are created in memory for speed. A set of disk-writing threads, working in parallel with the stream-receiving and flat-field-correcting threads, learns via a standard mutex signal of any tiff generations, and goes on to write the files to a 1.7 PB BeeGFS storage (task (f)).

The default ARAPAHO configuration, as used at the P14 beamline, includes 176 stream-receiving and processing threads, 44 disk-writing threads, and 3600 memory buffers each big enough to hold a final tiff. ARAPAHO runs on a 88-core DELL R830 server, hyper-threaded for a total of 176 hyper-threads, with 256 GB RAM.

The PCO CAMERA server is by default configured with 1 data-extraction thread, 2 sending threads, and 1800 memory buffers each big enough to hold a raw frame. It is running on a 20-core/40-hyper-thread server equipped with 96 GB RAM.

The data acquisition can be followed live via two separate viewers running in parallel, one for the raw data, and one for the flat-field corrected frames. For the raw data we use ADXV [https://www.scripps.edu/tainer/arvai/adxv/adxv_1.9.10/AdxvUserManual_v1.1.pdf], which reads tiff files from BeeGFS and displays them on IMAGE TRACKER server (Nikolova, Bourenkov *et al*., manuscript in preparation) requests at the rate of 25 frames per second. Each tiff file is 8.1 MB. The flat-field-corrected frames are displayed by an in-house developed Python viewer, FFC VIEWER, at the rate at which ARAPAHO is sending them, 23 frames per second (Supplementary Material S1). The size of each flat-field-corrected frame is variable and dependent on the jpeg compression rate. It is currently limited to 2.6 MB. To date this has been sufficient for all samples imaged. During a user experiment the two viewers usually occupy a screen with a 4K resolution sharing it between themselves. The relatively high display rate achieved is attributed to an 18-core/36-hyper-thread user desktop computer, its careful kernel tuning, including increasing the default receiving network buffers, and the minimum RAM to be kept free, disabling swap, etc. (table, supplementary material). Additional factors playing a role in boosting the display rates are the very fast BeeGFS system in the case of ADXV, and the recent switch from UDP to TCP/IP in the case of the FFC VIEWER together with kernel tuning also on the data sending servers side.

After a successful data collection consisting of acquisitions at 4 distances the 3D reconstruction pipeline is automatically triggered. The pipeline consists of an application written in-house, TOMO CTF (C++), and a tomographic reconstruction using the “GridRec” algorithm embedded in TomoPy (Gürsoy *et al*., 2014). Due to the high-frequency beam oscillations affecting the background but not the object image, traditional flat-fielding (averaging all flat-field projections) tends to lead to suboptimal results. Therefore, only the three best matching background frames are selected instead – based on the correlations stored by ARAPAHO in the tiff header – and their average is used as the denominator during the flat-fielding of a projection frame. After correcting for lateral and vertical shifts that have occurred with the camera movement between the 4 acquisitions, the phase retrieval is carried out using the multi-distance contrast-transfer-function method (Cloetens, Ludwig, Baruchel, Van Dyck *et al*., 1999; Zabler *et al*., 2005). Subsequently, as the last step in TOMO CTF, the position of the rotation axis is determined. For both the shift and the rotation axis finding, image registration is performed via a Fourier-based correlation with a high-pass filter. By default we zero out 20 x 20 unique Fourier coefficients around the origin – analogous to excluding all image features bigger than about 100 pixels. We exclude in addition any feature that appears in one dimension only, zeroing out the Fourier coefficients along the origin row and column. With those defaults the results are accurate within a pixel for almost all samples and experimental conditions.

TOMO CTF is executed via slurm (“simple linux utility for resource management”, https://slurm.schedmd.com) on three cluster nodes in parallel utilising 264 cores simultaneously. This results in a CTF computing time of 20 seconds per dataset. TomoPy GridRec is likewise parallelized on three cluster nodes and completes in a total of 60 seconds. Both the CTF-corrected images, and the reconstructions are saved to the BeeGFS file system and can be inspected at the beamline right after the data acquisition using an in-house developed viewer, TOMO VIEWER, written in C++.

Scanning parameters, preview images and several tomogram slices for each sample are uploaded to the ISPyB database (Delagenière *et al*., 2011) automatically. Users and collaborators can review those online remotely via a web browser. Both the raw data, and the reconstructed data sets can be downloaded directly from the file system using an SSH File Transfer Protocol (SFTP) proxy connection.

### 2.5. Ethical Statement

No animals were sacrificed for the purpose of the presented study here. The analysed murine organs were harvested as a by-product of animal studies (heart, kidney, liver, brain: 33.33-42502-04-21/3668, lung: 33.19-42502-04-17/2630) approved by the ethics administration of Lower Saxony, Germany and performed in compliance with the guidelines of the German ethics law.

## 3. Results

### 3.1. High Throughput Tomography - HiTT

High Throughput Tomography (HiTT) combines fast data acquisition with streamlined data processing and 3D reconstruction. While 3rd and 4th generation synchrotron light sources enable rapid data acquisition because of their high photon flux, this also comes with a significant consequence - huge volumes of raw X-ray projection data are generated which need to be processed. This can be a bottleneck, especially for non-expert users of X-ray imaging facilities and beamlines. Therefore we aim to provide a complete solution that integrates and automates the data processing. Fig. 3 depicts the current HiTT pipeline. Phase contrast nano-holotomography is performed by acquiring X-ray projection datasets at four different propagation distances. Each data collection takes about 34 seconds. This includes moving the detector stage to the desired propagation distance, opening the fast X-ray beam shutter, acquiring 100 background flat-field frames and 1810 projection frames (1800 for reconstruction plus 10 additional frames for determining the centre of the rotation axis). This results in a total acquisition time of 2 min 16 s for all four distances. Subsequently, the data processing is triggered automatically. To this end the data processing is divided into two different steps. First a custom made C++-based software performs flat-field corrections and the contrast-transfer function phase retrieval. This step takes around 20 s of computation time and is followed by the Python-based 3D reconstruction step which takes around 1 min. More details about the data processing can be found in Section 2.4.

**Figure 3.**
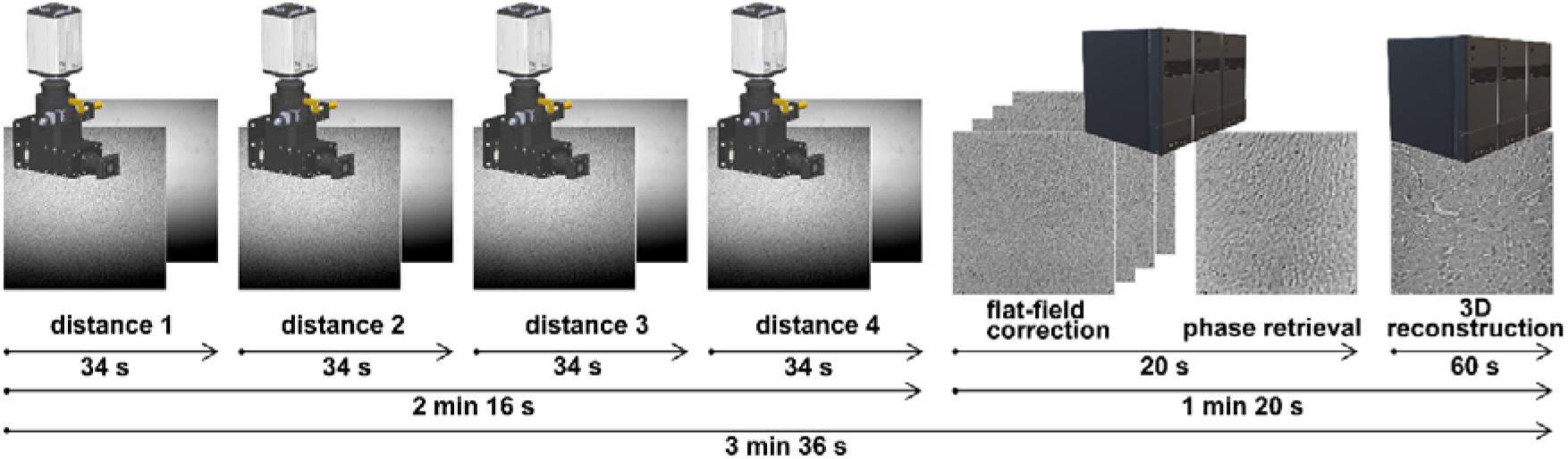
The HiTT pipeline. For each sample four different data collections at increasing propagation distances are performed. Subsequently the data processing is triggered automatically.

The MARVIN sample changer deployed at the P14 beamline is helping to increase the sample throughput as it removes the need for manual sample changes and therefore the breaking of the beamline hutch interlock system. Mounting a new sample takes 50 s until the scanning position is reached. If a sample is already mounted the exchange to a new sample takes 75 s. Sample mounting/dismounting can be performed in parallel with the data processing pipeline.

This combination of a sample changing robot and fast data acquisition leads to a total processing time of ca. 4 min and 47 s, where all data is processed simultaneously. The only manual input that is necessary is the positioning and centring of the sample inside the FOV. This is done by using the optical microscope embedded in the OAV system and the so-called “three click cantering” procedure which allows precise sample centring in a few seconds without the need for X-rays thus there is no “wasted” X-ray dose as the only X-ray exposure of the sample is for imaging. Under the generous assumption of 1 min for sample positioning a complete HiTT sample acquisition is performed in under 6 mins. Extrapolated this leads to a sample throughput of ∼ 10 samples/h or 80 samples per 8h beamtime shift. This includes both the raw imaging data and the 3D volume reconstructions. Thus the user leaves the facility with data that they can then interrogate at their leisure with their preferred 3D viewing and annotating software.

### 3.2. Flexibility of HiTT demonstrated on various sample types

In combination with the high-throughput and automation we are focussing on delivering a flexible X-ray imaging setup that is capable of acquiring high-quality 3D datasets from a wide variety of samples from different life-science disciplines including cells, organoids, unstained soft tissue samples and heavy-metal stained samples that are prepared for volume electron microscopy. The P14 beamline is easily tuneable between 6 and 30 keV using the double-crystal monochromator. For X-ray imaging we have successfully used an energy range of 10-27 keV. While being able to fine tune the acquisition parameters such as photon energy, exposure time or propagation distance comes with great flexibility, it also comes with the drawback that finding the right parameters can be tedious and time consuming which can be problematic given limited synchrotron beamtimes. Therefore, we have also been aiming at finding a sweet spot in imaging parameters that can be used for a broad range of samples.

Figure 4 shows a multitude of different types of samples that were all scanned with the following acquisition parameters: 4x1810 projection images with an exposure time of 10 ms, propagation distances of 61, 65, 71, 80 mm and 12.7 keV photon energy.

**Figure 4.**
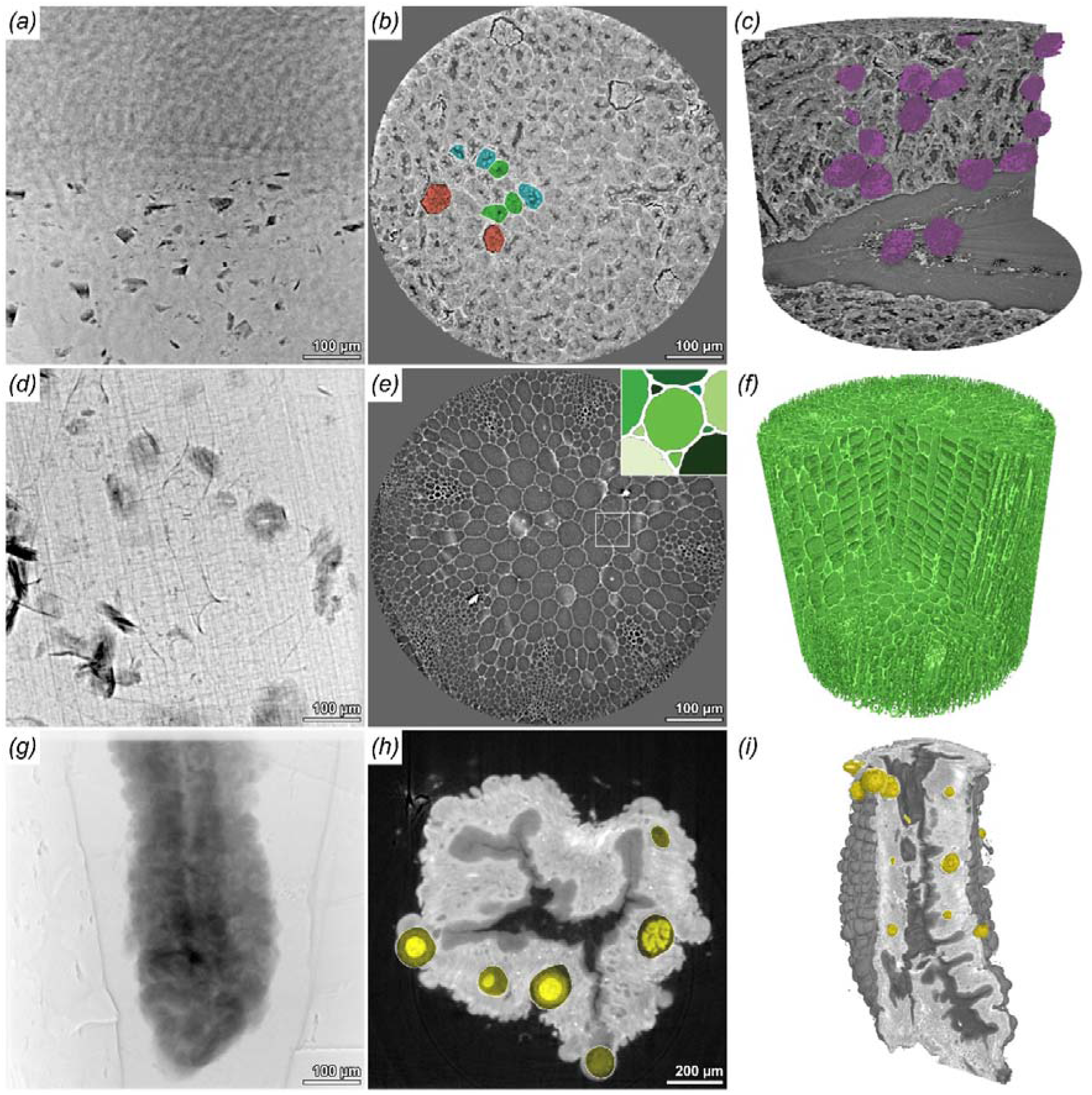
HiTT imaging can be successfully performed on various sample types. (a)-(c) projection image, reconstructed slice and 3D rendering of a paraffin embedded 1 mm murine kidney punch biopsy. Anatomical structures like glomeruli (red), distal convoluted tubules (cyan) and proximal convoluted tubules (green) can be easily discerned. (d)-(f) projection image, reconstructed slice and 3D rendering of a Arabidopsis thaliana stem. Zoom in in (e) shows that individual cells can be easily segmented and labelled. (g)-(i) projection image, reconstructed slice and 3D rendering of a osmium stained and resin embedded midgut of an Anopheles stephensi mosquito, which was infected with Plasmodium berghei. Oocysts formed by the parasites are clearly visible in (h, yellow) and segmented in (i).

A paraffin embedded murine kidney is depicted in Fig. 4a-c. Anatomical structures of the kidney cortex such as glomeruli and the convoluted tubules can be easily identified. The glomeruli, the spherical filtration units, were segmented and visualised. Based on this a clear application would be to use HiTT to investigate gross morphological differences within tissue for various animal disease models. With the rate of both data acquisition and analysis it is possible to study many samples very quickly allowing statistical differences to be analysed alongside the more visual differences discernible within the 3D volumes.

Studying the 3D architecture of plants is another successful example application for HiTT. Here we depict the stem of a 11 days old *Arabidopsis thaliana* seedling (Fig. 4g-i), cell walls give a large phase contrast signal which makes the segmentation of single cells easily feasible. Fig. 4h shows an example of a simple threshold segmentation with subsequent labelling of cells in different shades of green. Both samples discussed were prepared and scanned without any contrast enhancing agents and in different embedding media (paraffin, ethanol), but all show sufficient to very good contrast. Nevertheless HiTT can also be performed on heavy-metal stained samples, which are commonly used for lab-based μCT or (volume) electron microscopy (vEM). Fig. 4j-l displays an osmium stained, resin embedded midgut of an *Anopheles stephensi* mosquito, which was prepared for serial-blockface electron microscopy imaging. Due to the long scanning times in vEM (>days) acquiring an overview data set of the samples using X-ray tomography is of great benefit. For this example in the field of infection biology HiTT has been used to image malaria parasites within mosquito guts. The resolution was such that the oocysts (structures where malaria parasites mature) could be easily visualised and segmented (Fig. 4k,l). The imaging resolution is such that oocysts of different sizes could be detected, regardless of their distribution at the periphery of or inside the mosquito host mid-gut tissue. Given the speed with which data can be obtained it is possible to image many mosquito guts and build up a database or atlas of parasite presentation within the gut.

### 3.3. Using HiTT to perform virtual 3D histology

The 3D analysis of preclinical and clinical soft-tissue samples has become a great application for high-resolution x-ray tomography in recent years. The main disadvantage of conventional histology is its 2D nature, while “X-ray based virtual” offers non-destructive isotropic 3D imaging of tissue samples without interfering with standard (pre)-clinical histology protocols. Fig. 5 shows a collection of murine tissue biopsy samples displaying the capabilities of our HiTT setup for performing X-ray based virtual histology. We can demonstrate that using HiTT a resolution comparable to conventional histology can be achieved depicting the nuclei of neuronal cells (a,b), single lung alveoli (c), as well as striations and intercalated discs of cardiomyocytes in heart tissue (d). In medullary kidney tissue the fine tubular filtration network is displayed even including single erythrocytes in the blood vessels (e). Liver tissue organisation with a central vein of a hepatic lobule is displayed in (f).

**Figure 5.**
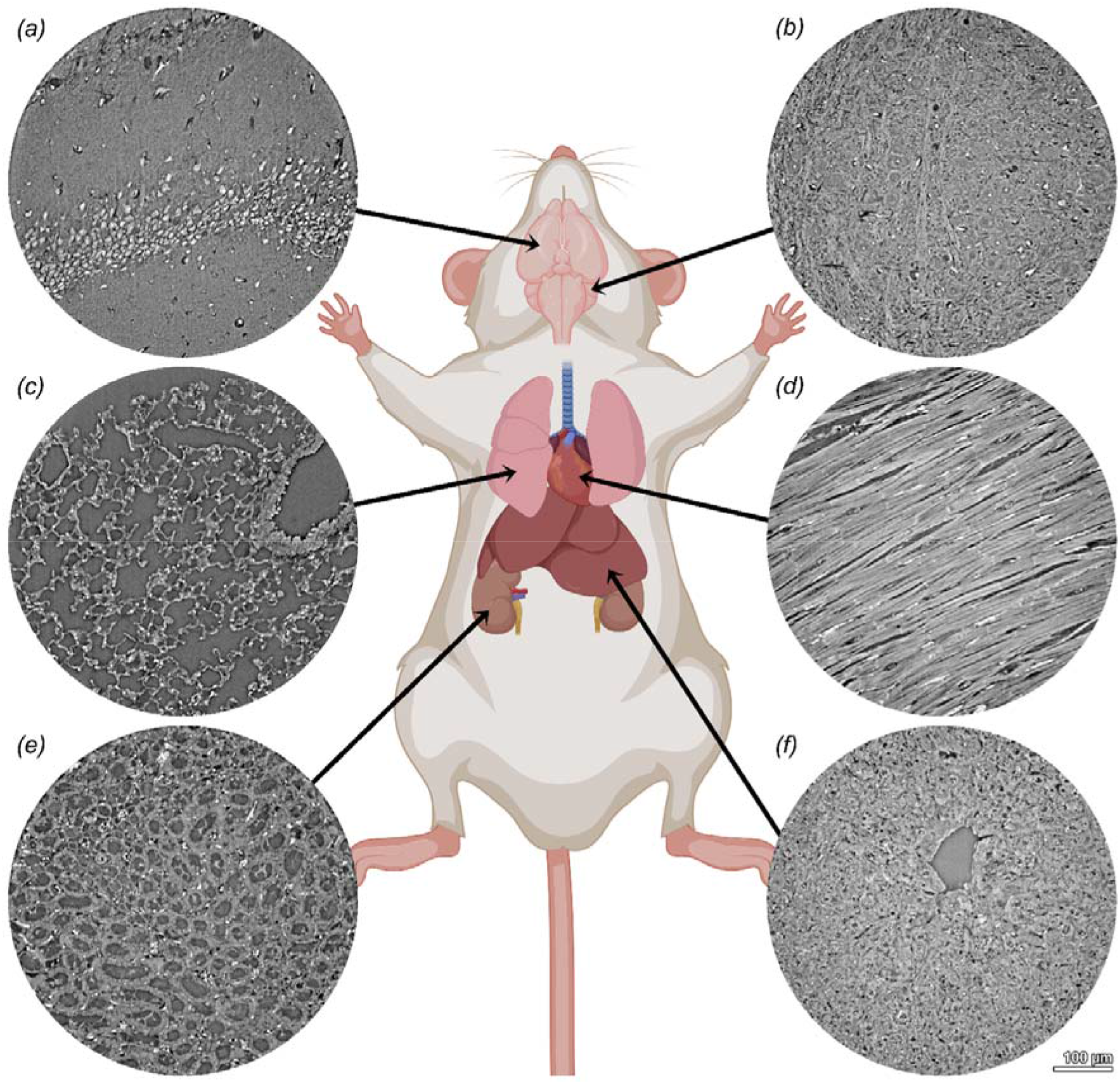
HiTT for virtual histology of murine tissues. One millimetre punch biopsies were taken from formalin fixed and paraffin embedded mouse organs and imaged at the P14 beamline. (a) brain (cerebrum), (b) brain (cerebellum), (c) lung, (d) heart, (e) kidney medulla, (f) liver. Mouse sketch was created with BioRender.com.

To understand pathologies in tissue samples it is often not enough to look at small regions, because the context of the (micro)-anatomical localisation of a structure or pathological alteration is important. High resolution data sets come in most applications with the caveat of a relatively small FOV which makes it difficult to put the acquired data into its tissue context. Therefore, we established a pipeline to perform tiled acquisitions in each dimension to be able to increase the FOV and to enable the scanning of larger samples. Due to the use of hard X-ray radiation larger samples (even in the cm range) can be penetrated easily and local tomography data collection can be performed. Fig. 6 shows a murine kidney punch biopsy of a formalin fixed paraffin embedded sample. Fig. 6a shows a single reconstructed and stitched slice, while Fig. 6b shows a false coloured 3D rendering of the same dataset. The data set consists of 20 single HiTT acquisitions (2x2x5) and has a FOV of 0.85x0.85x1.85 mm^3^. Here the differences in kidney architecture are visualised going from the kidney cortex with glomeruli and convoluted tubules (top) to the medulla (bottom). The 3D rendering displayed in Fig. 6b resembles the appearance of an haematoxylin and eosin staining displaying the similar appearance of HiTT virtual histology and conventional 2D histology. Videos of both representations can be found in the supplementary materials.

**Figure 6.**
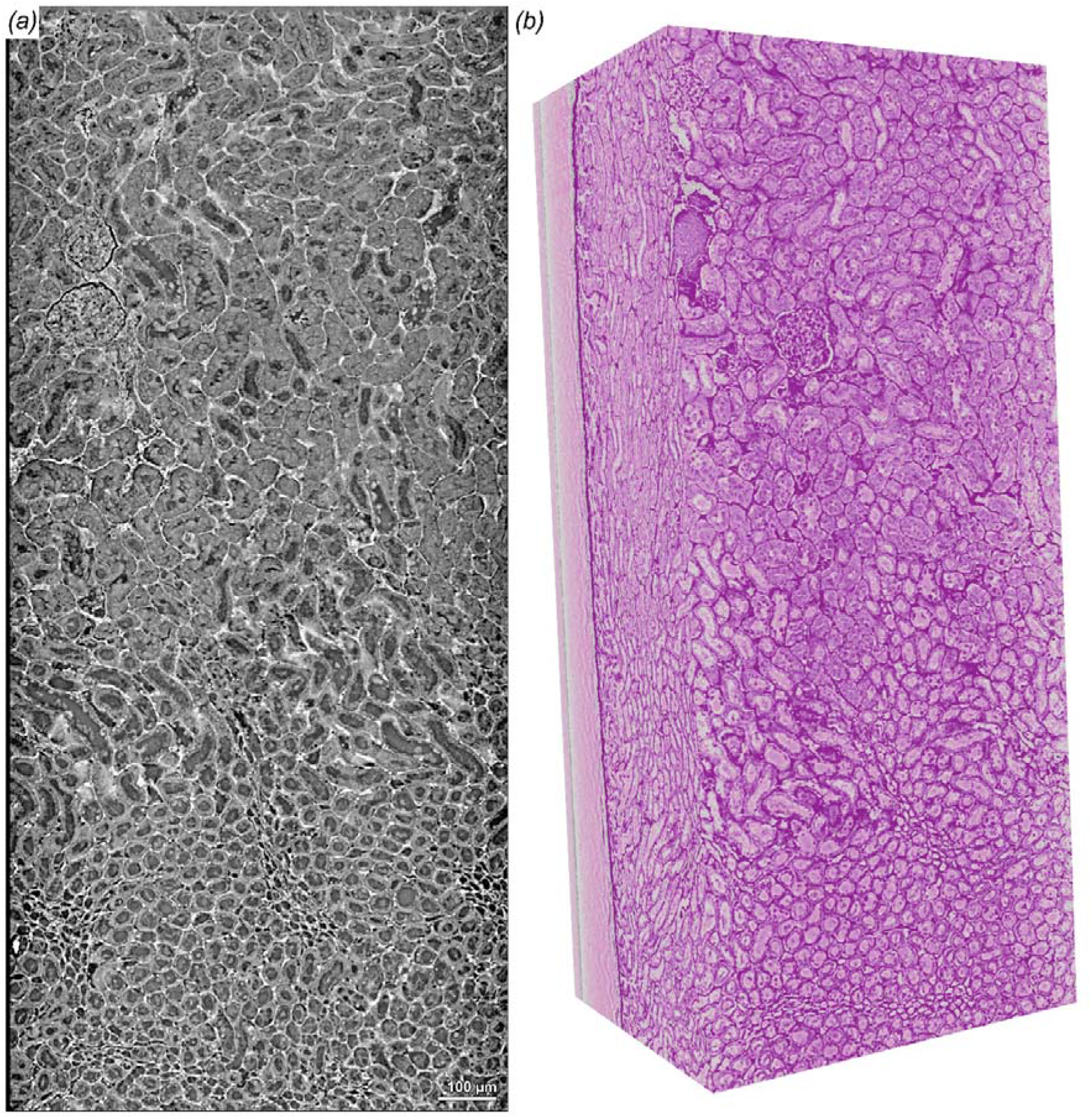
Stitched acquisition of a murine kidney punch biopsy. Twenty single acquisitions (2x2x5 scans) of a murine kidney punch biopsy were acquired and stitched together. (a) shows a single reconstructed slice displaying kidney tissue from the kidney cortex (top) to the medulla (bottom). (b) pseudo haematoxylin and eosin coloured 3D rendering of the same data set showing remarkable resemblance with conventional histology.

## 4. Conclusions and Future Prospects

We have presented high throughput tomography (HiTT) as a 3D X-ray imaging method capable of imaging soft biological samples of the order of 1mm in size at a cellular resolution. Taking advantage of the infrastructure established for macromolecular crystallography over the past decades, primarily the automated sample changer and extremely reliable beamline instrumentation controlled by an easy-to-use graphical user interface (GUI) enables the high throughput with very low sample to sample exchange time. Automatic triggering of the 3D reconstruction pipeline along with access to a highly optimised compute cluster results in 3D volumes available for interpretation within minutes of data collection completion. Whilst this approach is now taken for granted in synchrotron based macromolecular crystallography in our experience this is not the norm for X-ray imaging. Our aim with HiTT is to open up the field to biologists with minimal expertise in “hard core” X-ray imaging but who wish to see their samples in 3D and recognise that X-ray imaging will allow them to do this. We believe that this will be a game changer for biology. Going forward we plan to offer HiTT as a user facility at the EMBL Beamline P14 on Petra III alongside our current structural biology program (https://www.embl.org/groups/macromolecular-crystallography/).

## Acknowledgements

The authors thank the Instrumentation team, the IT team of EMBL Hamburg and Michael Agthe for technical support. In addition, we would like to thank Maxim Polikarpov and Ivars Karpics for their work in establishing the initial imaging setup P14. We thank Christian Dullin for providing murine tissue samples, Steffen Ostendorp for providing Arabidopsis samples and Friedrich Frischknecht for providing the *Plasmodium berghei* infected *Anopheles stephensi* mosquitos. Resin embedded mosquito tissues were prepared with the help of the Electron Microscopy Core Facility at EMBL Heidelberg. Access to the beamline is provided via the EMBL Hamburg User Program.

## Funding information

Nedal Darif is funded by the Deutsche Forschungsgemeinschaft (DFG, German Research Foundation) – Projectnumber 240245660 – SFB 1129 (project Z2).

## Notes

### Competing Interest Statement

The authors have declared no competing interest.

